# ChEC-seq produces robust and specific maps of transcriptional regulators

**DOI:** 10.1101/2021.02.11.430831

**Authors:** Gabriel E. Zentner, Robert A. Policastro, Steven Henikoff

## Abstract

We previously introduced chromatin endogenous cleavage and high-throughput sequencing (ChEC-seq), which uses fusion of a chromatin-binding protein of interest to micrococcal nuclease (MNase) and high-throughput sequencing to generate genome-wide maps of factor binding. Here, we respond to concerns that have been raised about the specificity of the method relative to its negative control when a single long calcium incubation time is used. Through reanalysis of our previously published datasets, we show that short-duration ChEC-seq experiments provide robust, specific maps of transcriptional regulators across the budding yeast genome. Our analyses also confirm that consideration of MNase digestion kinetics is important for proper design and interpretation of ChEC-seq experiments.

---

Pugh and colleagues report little difference between signals for MNase-tagged proteins and free MNase (that is, nuclear-localized MNase not fused to any factor) when a single long time point (5 minutes after calcium addition) is analyzed (Mittal et al., 2021). Enrichment is thus considered to be largely nonspecific. We originally performed a broad time course analysis of cleavage by the MNase-tagged general regulatory factors (GRFs) Abf1, Rap1, and Reb1 and free MNase. This revealed two general classes of binding sites: ‘fast’ sites, where maximal cleavage was reached ≤1 minute after calcium addition, and ‘slow’ sites, where cleavage increased up to 20 minutes after calcium addition. Thus, consideration of MNase cleavage kinetics is important for proper interpretation of ChEC-seq results.

Expression of free MNase in cells followed by calcium incubation is akin to the use of exogenous MNase in a standard epigenome profiling experiment, with diffusion, chromatin association, and cleavage kinetics unconstrained by fusion to a protein with affinity for specific loci. Thus, longer digestion times yield increased cleavage within accessible DNA, resulting in increased ‘nonspecific’ signal that negates the remaining specific signal. To illustrate this, we plotted free MNase signal around the TSSs of 4,857 genes encoding verified ORFs across a 40-fold range of calcium incubation times. We observed weak signal at 30 seconds and 1 minute of incubation with calcium (Fig. 1A-B), but by 5 minutes noted robust cleavage within promoter NDRs and nucleosome linkers, leading to an uneven background.

**Figure 1.**
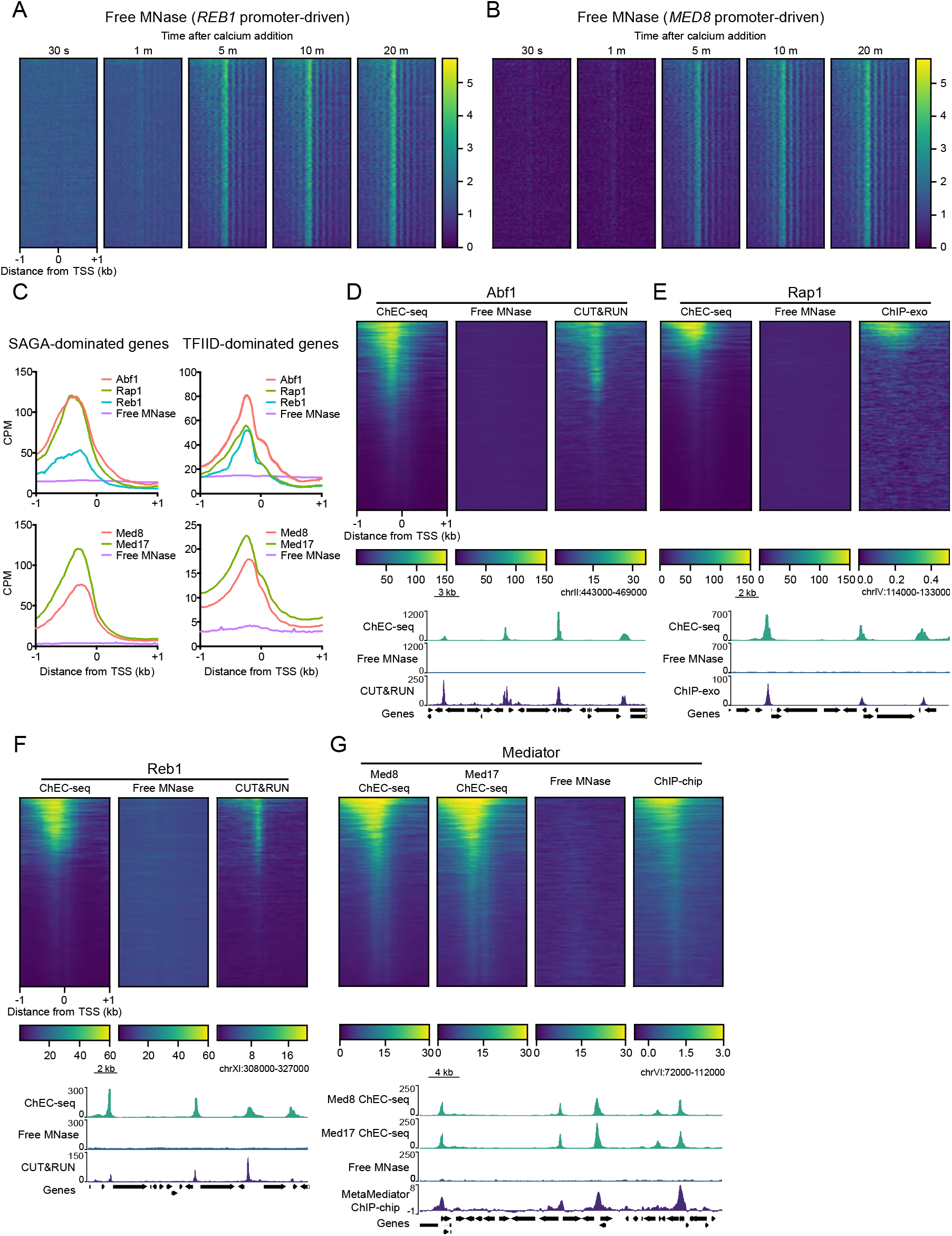
Short-duration ChEC-seq experiments show high specificity. Heatmaps of CPM-normalized free MNase signal around 4,857 TSSs. Each heatmap is independently sorted descending by the average signal in the 2 kb window surrounding each TSS. (C) Average plots of CPM-normalized GRF (merged 10 s – 1 m time points), Mediator subunit (merged 30 s and 1 m time points), and free MNase (merged 10 s – 1 m time points for W1588-4C, merged 30 s and 1 m time points for BY4705) signal at SAGA-dominated and TFIID-dominated genes. (D) Heatmaps of pooled 10 s – 1 m Abf1 and W1588-4C free MNase ChEC-seq and pooled 1 s – 128 s Abf1 CUT&RUN at 4,857 TSSs. All heatmaps are sorted descending by average Abf1 ChEC-seq signal in the 2 kb window surrounding each TSS. Also shown are genome browser tracks of the same data along a representative segment of the yeast genome. (E) Same as (D) but for Rap1 and with pooled Rap1 ChIP-exo signal. (F) Same as (D) but for Reb1. (G) Heatmaps of pooled 30 s and 1 m Med8, Med17, and BY4705 free MNase ChEC-seq and MetaMediator (combined ChIP-chip signal for 12 Mediator subunits) data from WT yeast. All heatmaps are sorted descending by average Med8 ChEC-seq signal in the 2 kb window surrounding each TSS. Also shown are genome browser-style tracks of the same data along a representative segment of the yeast genome. See Fig. 2 for genome browser-style views of all analyzed datasets at each region shown here.

Having established that short calcium incubations produce relatively uniform background cleavages by free MNase, we assessed the binding of GRFs and Mediator at genes classified as dominated by the SAGA or TFIID coactivator complexes in their regulation (Huisinga and Pugh, 2004) using merged ≤1 minute calcium incubation time points for our previously published factor-specific and free MNase ChEC-seq datasets (Grünberg et al., 2016; Zentner et al., 2015). At both gene sets, we detected robust enrichment of all factors over free MNase on average (Fig. 1C).

We initially reported numbers of fast GRF binding sites lower than the approximate number of RNA polymerase II (RNAPII)-transcribed genes in the yeast genome (Abf1, 3,702; Rap1, 1,974; Reb1, 2,711) (Zentner et al., 2015) and did not annotate the genes associated with these sites. Thus, it was unclear how many genes are associated with rapid, specific cleavage by GRF-MNase fusions. To assess this, we have visualized GRF ChEC-seq and free MNase signal around TSSs as heatmaps. Approximately one-third of the analyzed TSSs displayed upstream enrichment of Abf1 by ChEC-seq and CUT&RUN (Skene and Henikoff, 2017) (Fig. 1D). Enrichment of Rap1 was observed at ~15% of the analyzed TSSs by both ChEC-seq and ChIP-exo (Rossi et al., 2018) (Fig. 1E). We found robust Reb1 enrichment at ~25% of analyzed TSSs by ChEC-seq and CUT&RUN (Fig. 1F). We similarly assessed Mediator enrichment around TSSs. Approximately 25% of TSSs displayed strong upstream enrichment of both Med8 and Med17; however, modest signal was observed for most of the remaining TSSs relative to free MNase (Fig. 1G). This pattern was also seen in no-tag-normalized Mediator ChIP-chip data (Jeronimo et al., 2016). We conclude that ChEC-seq indeed reveals widespread binding of Mediator to the upstream regions of most RNAPII-transcribed genes, consistent with the global negative effect of Mediator dissociation on RNAPII occupancy (Warfield et al., 2017). Visualization of all datasets at the loci examined in Fig. 1D-G also revealed common and distinct peaks (Fig. 2), supporting capture of specific binding events by short-duration ChEC-seq experiments.

**Figure 2.**
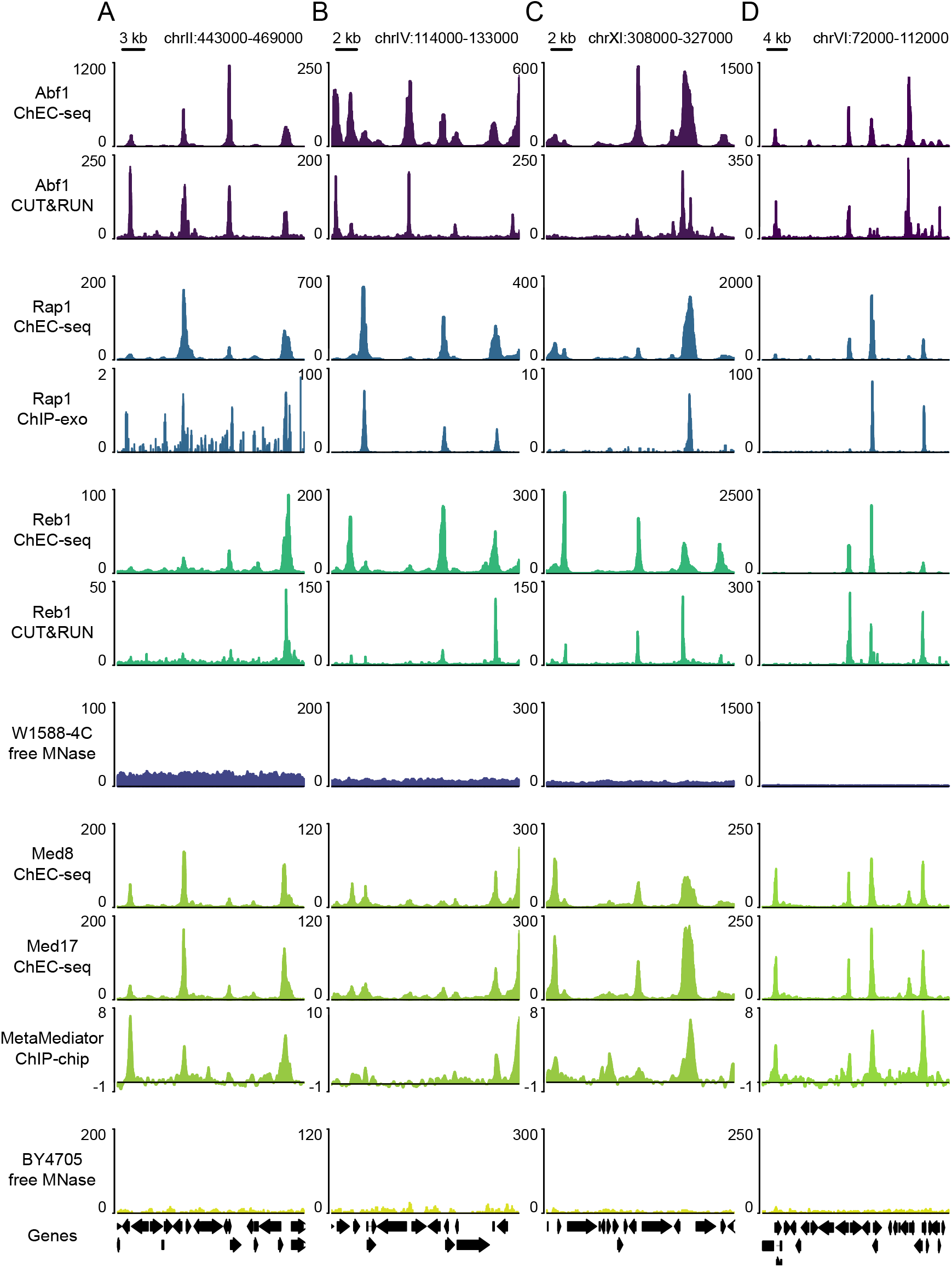
Landscapes of GRF and Mediator binding to the yeast genome. Genome browser-style visualization of all analyzed ChEC-seq, CUT&RUN, and ChIP-exo datasets at the four regions of the yeast genome displayed in the analysis of (A) Abf1, (B) Rap1, (C) Reb1, and (D) Mediator ChEC-seq data presented in Fig. 1D-G.

We next called peaks of GRF ChEC-seq binding and assessed motif enrichment. We observed bimodal enrichment of known motifs around peak summits, consistent with cleavage of DNA adjacent to the motif, and detected robust consensus sequences in peaks *de novo* (Fig. 3A-C). Short-duration ChEC-seq experiments thus capture sequence-specific binding of GRFs, consistent with our detection of strong motifs in ‘fast’ sites (Zentner et al., 2015). We also determined the fraction of reads in peaks (FRiP) (Landt et al., 2012) for each dataset after down-sampling and peak calling. All three GRF ChEC-seq datasets displayed exceptional signal-to-noise ratios, with FRiP values of 0.45-0.65 obtained using just ~25,000 reads for peak calling (Fig. 3D). Numbers of called peaks increased with additional reads considered and plateaued at ~2 million reads with FRiP values of 0.69-0.84, indicating that at least two-thirds of all reads from each dataset is contained within peaks called using ~2 million reads. Together with the above analyses of motif enrichment, these results confirm that short-duration ChEC-seq produces high-quality data for GRFs.

**Figure 3.**
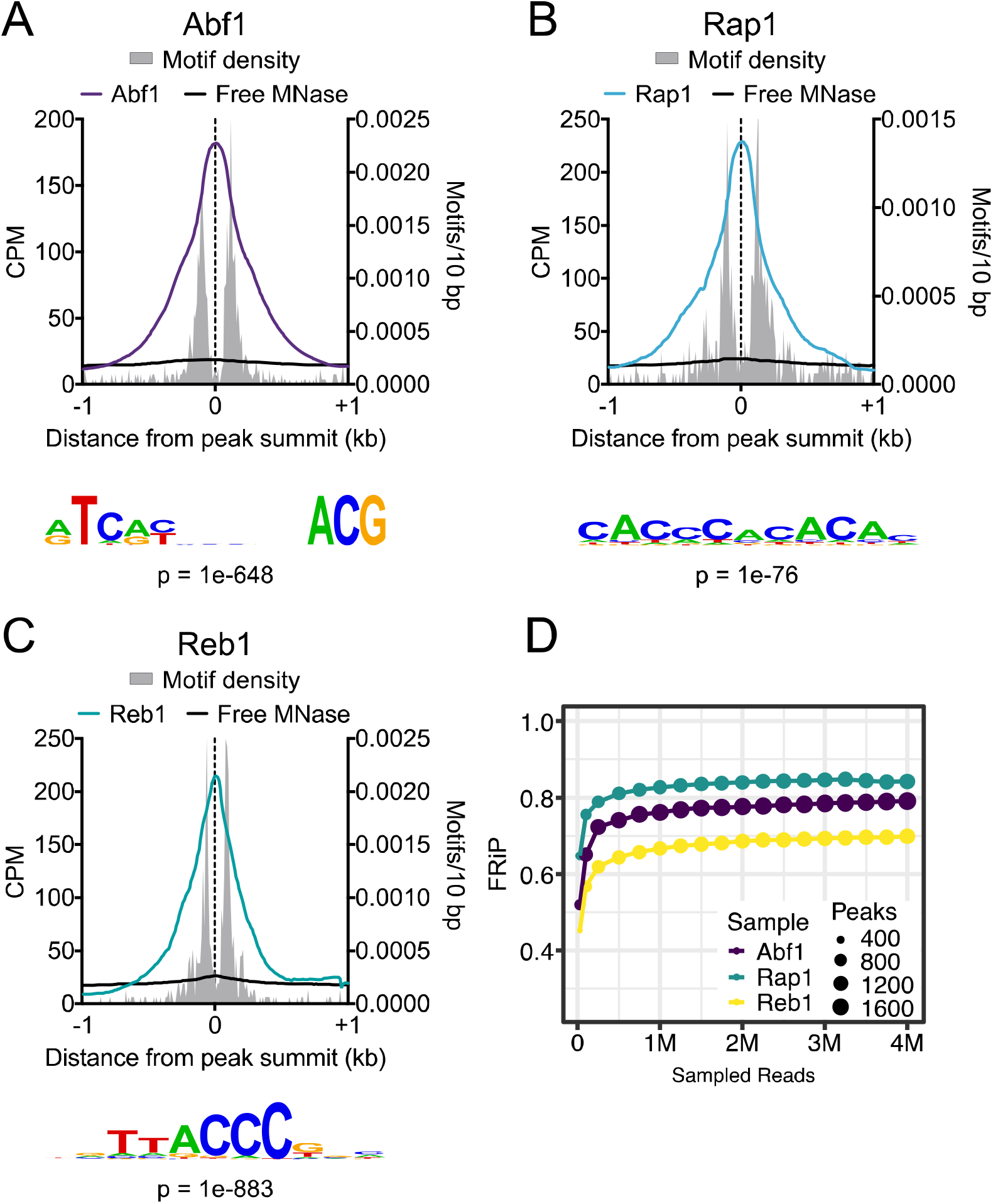
ChEC-seq detects sequence-specific binding of GRFs. Average plots showing ChEC-seq signal and motif enrichment in 10 bp bins around the summits of (A) 2,098 Abf1 peaks, (B) 1,623 Rap1 peaks, and (C) 1,564 Reb1 peaks. The sequence logo of the top motif match discovered *de novo* with peaks for the indicated factor is shown below each plot. (D) Fraction of reads in peaks (FRiP) values and peak numbers for down-sampled GRF ChEC-seq datasets. Down-sampled read numbers were 25,000, 100,000, and then every 250,000 starting at 250,000.

We have shown that short-duration calcium incubations yield robust and specific signal for GRFs and Mediator as measured by enrichment over free MNase, signal from orthogonal methods, and the presence of consensus motifs (for GRFs). The apparent lack of specificity in ChEC-seq experiments in which long calcium incubation is used is likely due to two additive factors – decreased specific signal in the ChEC-seq experiment and increased nonspecific signal in the free MNase experiment. Detection of these factors was made possible by the use of time courses for both factor-specific and free MNase experiments, as performed in the original ChEC-seq paper (Zentner et al., 2015) and recommended in a related methods article (Grünberg and Zentner, 2017). Given the increase in free MNase signal at longer time points, comparison of short-duration MNase fusion and free MNase experiments (≤1 minute) yields the most robust results. Similar observations were made in a recent study that used ChEC-seq to characterize the contribution of intrinsically disordered regions of transcription factors to their DNA-binding activities (Brodsky et al., 2020). The authors performed a calcium digestion time series and chose short incubation times (30-60 sec) for their experiments.

As a complement to this article, we provide a detailed description of our current best practices for ChEC-seq at protocols.io (https://www.protocols.io/view/chromatin-endogenous-cleavage-and-high-throughput-bgthjwj6). We also draw attention to two additional responses to these critiques from the Hahn (Donczew et al, bioRxiv 2021) and Shore (Bruzzone et al, bioRxiv 2021) labs.

## Methods

### Datasets

FASTQ files were obtained from the SRA: GRF and W1588-4C free MNase ChEC-seq (SRP056746) (Zentner et al., 2015); Mediator and BY4705 free MNase ChEC-seq (SRP074777) (Grünberg et al., 2016); Abf1 and Reb1 CUT&RUN (SRP078609) (Skene and Henikoff, 2017); Rap1 ChIP-exo (SRP096827) (Rossi et al., 2018). The MetaMediator track was obtained from the supplemental of the original publication (Jeronimo et al., 2016) and converted to a bigWig file with UCSC *bedGraphToBigWig*.

### Read alignment and processing

Paired FASTQ files were aligned to the sacCer3 genome assembly with Bowtie2 (Langmead and Salzberg, 2012) using default parameters plus ‘--no-unal --no-mixed – no-discordant --dovetail -I 10 -X 700’. Alignment SAM files were converted to sorted and indexed BAM files with SAMtools (Li et al., 2009). BAM files were merged with SAMtools. For GRF ChEC-seq analysis, 10 s, 20 s, 30 s, 40 s, 50 s, and 1 m time points were merged. For Mediator ChEC-seq analysis, 30 s and 1 m time points were merged. For CUT&RUN analysis, 1 s, 2 s, 4 s, 8 s, 16 s, 32 s, 64 s, and 128 s time points were merged. For ChIP-exo analysis, four replicates of Rap1 ChIP-exo were merged. Unless otherwise specified, all downstream analyses were performed with components of deepTools2 (Ramírez et al., 2016).

### Read coverage

Read coverage bigWig files were generated with *bamCoverage* with a bin size of 1 and CPM normalization. For Figure 1A-B heatmaps, 25 bp reads were not extended, allowing sharper visualization of periodic gene body signal. Reads were otherwise extended using *bamCoverage* -e, which extends reads to their fragment length based on mate-pair distance. CUT&RUN coverage was computed using the additional flag ‘--maxFragmentLength 120’ to exclude nucleosomal fragments as described in the original CUT&RUN publication (Skene and Henikoff, 2017). Genome browser visualization was performed with Gviz (Hahne and Ivanek, 2016).

### Heatmaps and average plots

The list of 4,857 TSSs used was generated by intersecting a published TSS list (Xu et al., 2009) with the verified ORF list from the Saccharomyces Genome Database. This list was then intersected with a list of SAGA/TFIID classifications (Huisinga and Pugh, 2004), yielding TSSs of 436 SAGA-dominated and 4,052 TFIID-dominated genes. TSS lists were then used to generate matrices of signal with *computeMatrix*. Heatmaps were generated with *plotHeatmap*. Average plot values were obtained with *plotProfile* and plotted in GraphPad Prism 7.

### Peak calling

GRF peaks were called using MACS2 (Zhang et al., 2008) with the merged W1588-4C free MNase dataset as the control file and an effective genome size of 12,100,000. To calculate FRiP values, GRF ChEC-seq alignment BAM files were downsampled with SAMtools and peaks were called as above. Fragments in peaks were counted with the *featureCounts* tool of Rsubread (Liao et al., 2019) and the assignment status of each fragment was output in CORE format. A fragment was required to have at least 50 bp overlap with a peak to be assigned to that peak. The fraction of fragments assigned to a peak was considered to be the FRiP for that dataset. BAM downsampling, peak calling, and read assignment to peaks were automated using a custom conda environment with bash and R scripts (https://github.com/rpolicastro/chip_downsampling).

### Motif analysis

To determine motif enrichment around peak summits, we used HOMER (http://homer.ucsd.edu) (Heinz et al., 2010) *annotatePeaks.pl* in motif finding mode (‘-m’) with 10 bp bins and the corresponding recommended position frequency matrix from ScerTF (Spivak and Stormo, 2012) after conversion to HOMER format. Position frequency matrices were derived from the following publications: Abf1 (MacIsaac et al., 2006), Rap1 (Morozov and Siggia, 2007), Reb1 (Badis et al., 2008). *De novo* motif searching was performed with HOMER *findMotifsGenome.pl* with a 400 bp window around peak summits with 6,516 yeast promoters (−1000 to +100 bp relative to the TSS with chrM sequences removed) as background regions. Based on the widths of recommended consensus motifs from ScerTF, motif size ranges of 13-17 bp, 11-15 bp, and 7-11 bp for Abf1, Rap1, and Reb1, respectively, were used.

## Acknowledgements

We thank Jorja Henikoff for analytical assistance. This work was supported by NIH grants R35GM128631 (G.E.Z.) and R01HG010492 (S.H.) and the Howard Hughes Medical Institute (S.H.)

